# *In vivo* sequential mutagenesis in germinal center B cells using a dual-recombinase approach: FOXO1 re-expression upon FOXO1 knockout rescues class switch recombination

**DOI:** 10.1101/2024.05.17.592915

**Authors:** Carlota Farré Díaz, Eleni Kabrani, Wiebke Winkler, Claudia Salomon, F. Thomas Wunderlich, Martin Janz, Klaus Rajewsky

**Affiliations:** Immune Regulation and Cancer, Max Delbrück Center for Molecular Medicine in the Helmholtz Association, Berlin 13125, Germany; Biology of Malignant Lymphomas, Max Delbrück Center for Molecular Medicine in the Helmholtz Association, Berlin 13125, Germany; Experimental and Clinical Research Center, a cooperation between the Max Delbrück Center for Molecular Medicine in the Helmholtz Association and the Charité – Universitätsmedizin Berlin, Berlin 13125, Germany; Hematology, Oncology and Cancer Immunology, Charité – Universitätsmedizin Berlin, Berlin 13125, Germany; Genome Diversification & Integrity, Max Delbrück Center for Molecular Medicine in the Helmholtz Association, Berlin 13125, Germany; Faculty of Medicine and University Hospital Cologne, Center for Molecular Medicine Cologne (CMMC), University of Cologne, Cologne 50931, Germany; Cologne Excellence Cluster for Stress Responses in Ageing-Associated Diseases (CECAD), University of Cologne, Cologne 50931, Germany; Max-Planck-Institute for Metabolism Research, Cologne 50931, Germany

**Keywords:** germinal center, sequential mutagenesis, dual-recombinase, Cre, Dre

## Abstract

Modeling complex (patho)physiological processes by sequential mutagenesis in mice is limited by the lack of optimized genetic tools and complex breeding strategies. We present a new Cre/DreERT2 dual-recombinase germinal center B-cell (GCBC)- specific strain, with co-expression of the recombinases from a single allele. This enables highly efficient Cre-mediated FOXO1 knockout followed by time-controlled, efficient Dre-mediated FOXO1 re-expression and functional rescue in GCBCs, demonstrating suitability for precise targeted sequential mutagenesis *in vivo*.

---

Site-specific recombination systems, such as Cre/loxP, are key technologies to study gene function and (patho)physiological processes *in vivo*. Dual-recombinase approaches represent a further technical development that enables more precise disease modelling, in particular with respect to multistep genetic manipulations and lineage tracing^1,2^. However, one limitation of the currently available systems is that recombinases are expressed from different loci, hereby requiring the generation of complex compound mutant mice – containing separate transgenes encoding the recombinases together with the respective target alleles – and thus complicating the analysis both in terms of breeding time frame and fidelity of sequential mutagenesis at the level of individual cells. Here, we describe a dual-recombinase mouse strain in the context of germinal center (GC) B cells (GCBCs), where Cre and the tamoxifen (TAM)- dependent DreERT2 recombinases are co-expressed from the immunoglobulin heavy constant gamma 1 (*Ighg1/Cγ1*) locus and thus at the initiation of the GC response. By sequentially targeting the endogenous *Foxo1* locus during the GCBC reaction as a proof-of-concept experiment, we demonstrate highly efficient Cre-mediated FOXO1 knockout (KO) with perturbation of GC physiology, followed by time-controlled Dre-mediated FOXO1 re-expression and phenotypic rescue, demonstrating that this system can be used *in vivo* as a genetic recombination switch that is precisely controlled with respect to cellular compartment and time.

Upon antigen encounter, mature B cells undergo clonal expansion in distinct histological structures called GCs. During the early phases of GC formation, activated B cells diversify their immunoglobulin repertoire via class switch recombination (CSR), and later, in the course of the GC reaction, via somatic hypermutation (SHM) of antibody V region genes^3^. Positively selected GCBCs differentiate into memory B cells (MBCs) or antibody-secreting plasma cells (PCs), thus shaping the humoral immune response^3^. Both CSR and SHM require the introduction and the efficient repair of DNA strand breaks in the immunoglobulin loci^4^. Errors during these processes can lead to oncogenic mutations and/or translocations, rendering (post)-GCBCs the origin of most B cell malignancies^5^. In order to study the physiology and malignant transformation of GCBCs, a number of Cre mouse lines have been generated^6,7,8,9,10^. However, none of the currently available lines allows for the genetic manipulation of GCBC differentiation in a stepwise fashion. This technical limitation prevents not only the *in vivo* investigation of the role and cooperation of genes at different time points during the GC reaction, but also the study of the sequential acquisition of oncogenic events in this microenvironment, underlying malignant transformation.

Addressing this experimental challenge, we have generated a novel dual-recombinase GCBC-specific mouse strain, in which Cre and the TAM-inducible DreERT2 recombinases are concomitantly expressed from the *Ighg1/Cγ1* locus, hereafter called the *Cγ1-CDE* strain (Fig. 1A). *Cγ1* encodes the constant region of IgG1. Upon T cell-dependent immunization, most activated B cells that enter the GC produce germline ɣ1 transcripts in response to IL4^6^, allowing the expression of both recombinases – and thus Cre activity – from the *Cγ1* locus early on. Cells that have completed switch recombination to IgG1 continue to express the recombinases during the course of the GC reaction. Since DreERT2 activation is dependent on TAM administration, the second (Dre-mediated) recombination event can be induced in a time-controlled manner, selectively in cells having undergone Cre induction. To functionally validate the newly generated strain, *Cγ1-CDE* mice were crossed to the fluorescent reporter strains *Rosa 26 (R26)-BFP*^*stopF*11^ and *R26-ZsGreen*^*stopRox*12^ for detection of Cre- and Dre-mediated recombination, respectively. *Cγ1-CDE, R26-BFP*^*stopF*^, *R26-ZsGreen*^*stopRox*^ compound mutant mice were immunized with NP-CGG and TAM was administered at days 2-5, 9-12 or 15-18 after immunization with subsequent flow cytometric analysis at days 7, 14 or 21, respectively (Fig. 1B-G). Consistent with the previous characterization of the *Cγ1-Cre* allele^6^, up to 96% of splenic GCBCs were labelled through Cre-mediated recombination (BFP^+^ GCBCs, Fig. 1B-D, Supplemental Fig.1A-B). Further gating on these BFP^+^ GCBCs showed a robust and highly efficient Dre-mediated recombination at all time points analyzed (Fig. 1B-D), with the highest dual labelling efficiency on day 7 of analysis (up to 60% ZsGreen^+^ cells within the BFP^+^ GCBC population, Fig. 1B). GCs are organized into two phenotypically and functionally distinct compartments, the dark zone (DZ) and the light zone (LZ)^3^. GCBCs in both compartments were successfully labeled (Fig. 1E-G). Importantly, in the absence of TAM, no Dre recombination was detected, demonstrating that Dre-mediated recombination is tightly controlled *in vivo* (Fig. 1E-G).

**Figure 1.**
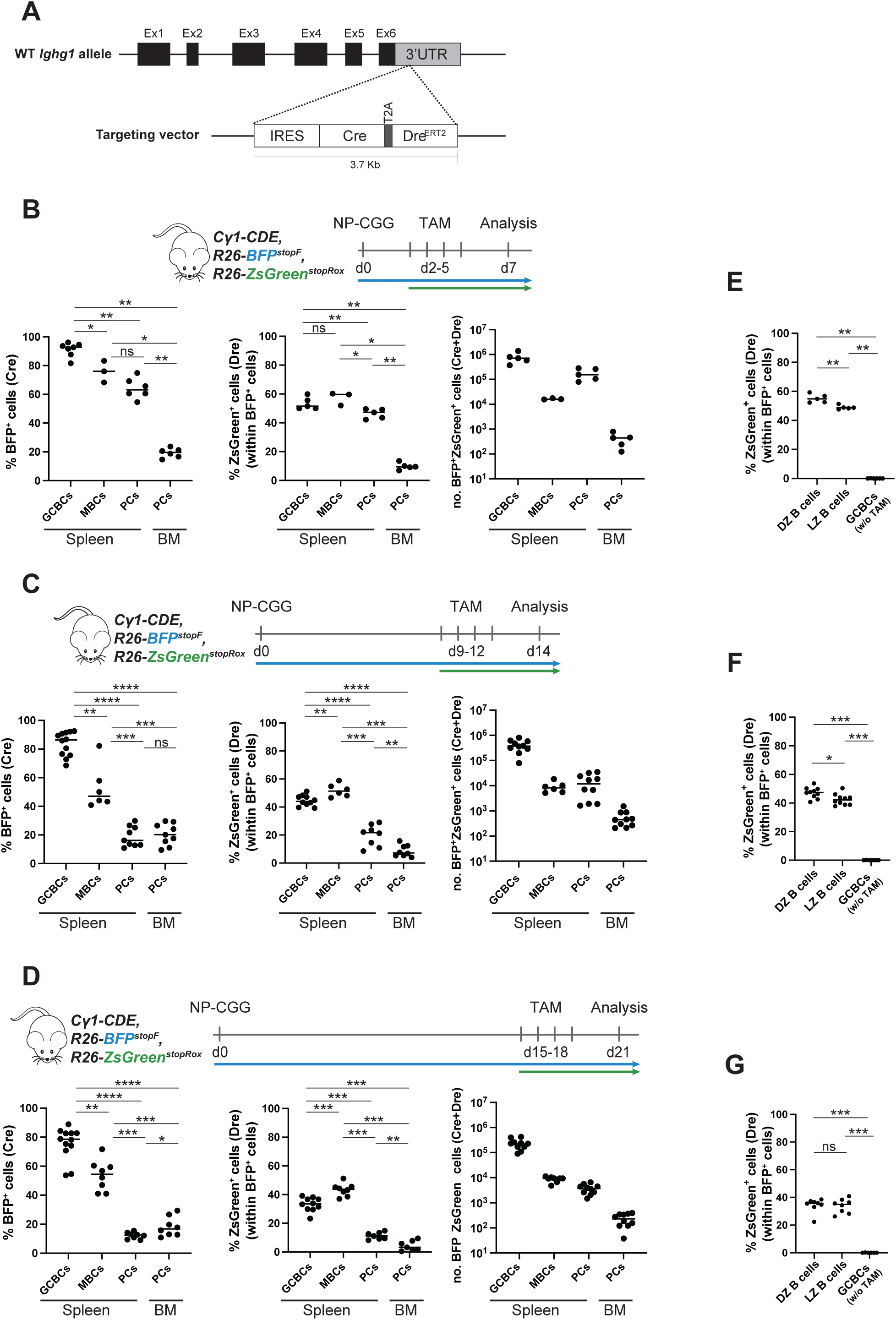
Characterization of the *Cγ1-CDE* strain. (A) Targeting strategy for the insertion of the IRES-Cre_T2A_DreERT2 cassette into the 3’-UTR of the mouse *Cγ1* locus. (B-G) Experimental scheme using the *Cγ1-CDE, R26-BFP*^*stopF*^, *R26-ZsGreen*^*stopRox*^ compound mutant mice, immunizing with NP-CGG at day 0, administrating TAM at day 2-5 (B and E), 9-12 (C and F) or 15-18 (D and G) and analyzing at day 7, 14 or 21, respectively. (B-D) Percentage of Cre-mediated BFP^+^ (left panel) and Dre-recombined ZsGreen^+^ (within the BFP^+^) cells (middle panel) among splenic Fas^+^CD38^-^ GCBCs, splenic Fas^-^CD38^+^IgD^-^IgG1^+^ MBCs and CD138^+^TACI^+^ PCs from spleen and BM (from one femur and tibia); as well as corresponding absolute numbers of the BFP^+^ZsGreen^+^ cells (right panel). (E-G) Percentage of ZsGreen^+^ (within BFP^+^) CXCR4^hi^DC86^low^ DZ and CXCR4^low^CD86^hi^ LZ GCBCs at day 7 (E), day 14 (F) and day 21 (G) after NP-CGG immunization. Data points for GCBCs without TAM administration are compiled from the three time points analyzed. Statistics: Mann-Whitney test \**P* ≤ 0.05; \*\**P* ≤ 0.01; \*\*\**P* ≤ 0.001; **** *P* ≤ 0.0001; ns, not significant. Each dot represents one mouse. Data from at least two independent experiments. Horizontal lines indicate the median.

Germline ɣ1 transcripts are detected in the activated B cell pool early on upon immunization, and their presence is independent of the subsequent IgG1 CSR event *per se*^13,6^. Accordingly, Cre-mediated followed by Dre-mediated sequential mutagenesis is achieved not only in IgG1^+^ (up to 70% ZsGreen^+^ within BFP^+^ cells) but also in IgM^+^ (up to 42% ZsGreen^+^ within BFP^+^ cells) GCBCs in our mouse model at all three time points analyzed (Supplemental Fig. 1E-F).

In addition, analysis of the post-GC compartments revealed the feasibility of labeling these cell types as well. MBCs exhibited solid Cre-mediated and Dre-mediated sequential mutagenesis (up to 84% BFP^+^ cells and 60% ZsGreen^+^ within BFP^+^ cells, Fig. 1B-D, Supplemental Fig. 1C). Analysis of the splenic plasmablast/PC compartment (Fig. 1B-D, Supplemental Fig. 1C) exhibited the highest labelling efficiency for this compartment at day 7 of analysis (up to 75% BFP^+^ cells and 50% ZsGreen^+^ within BFP^+^ cells, Fig. 1B), indicating the feasibility to target and thus study extrafollicular PCs with our *Cγ1-CDE* strain. However, the fraction of labelled PCs is reduced when analyzing the immunized mice at later time points (day 14 and day 21, Fig. 1C-D), possibly due to the short lifespan of extrafollicular PCs.

The FOXO1 transcription factor, although dispensable for GC development/maintenance, positively regulates both GC compartmentalization (DZ compartment) and CSR^14,15^. To this end, and as a proof-of-concept experiment, we aimed at genetically manipulating the endogenous *Foxo1* locus in a sequential manner. In addition to the already published Cre-inducible *Foxo1* null allele (*Foxo1*^*fl*^)^16^, we generated a rox-controlled, Dre-inducible *Foxo1* allele (*Foxo1*^*stopRox*^). The *Foxo1*^*stopRox*^ allele carries a STOP cassette flanked by rox sites in front of the endogenous *Foxo1* open reading frame, leading to the inactivation of the allele in the absence of Dre activity (Supplemental Fig. 2A). Of note, while hemizygous *Foxo1* mice are viable and fertile, homozygous germline deletion of FOXO1 results in embryonic lethality^17,18^. In order to verify that the *Foxo1*^*stopRox*^ allele is functional upon deletion of the STOP cassette, splenic B cells from *Cγ1-CDE, Foxo1*^*stopRox/wt*^ compound mutant mice were activated *in vitro* and cultured in the presence or absence of 4-hydroxytamoxifen (4-OHT), followed by sequencing analysis of *Foxo1* transcripts (Fig. 2A). Indeed, expression of the targeted allele was observed solely in the presence of 4-OHT, demonstrating not only that the *Foxo1*^*stopRox*^ allele is functional, but also the tightness of the system. *In vivo*, crossing of *Cγ1-CDE, Foxo1*^*stopRox*^ mice with *Foxo1*^*fl*^ transgenic mice (*Cγ1-CDE, Foxo1*^*fl/stopRox*^) would generate a scenario in which upon NP-CGG immunization, and therefore Cre activation, the *Foxo1*^*fl*^ allele would be deleted, resulting in a FOXO1-KO condition in GCBCs (Fig. 2B). GCBC-specific FOXO1 deletion is expected to lead to loss of the DZ compartment and to defects in CSR, while maintaining the GCBC phenotype^11,12^. In a second step upon TAM treatment – and hence Dre nuclear translocation and activation –, re-expression of FOXO1 would be achieved from the *Foxo1*^*stopRox*^ allele, thus leading to a rescue of the previously induced FOXO1 deficiency (Fig. 2B). To address whether the sequential deletion and re-activation of FOXO1 from the endogenous locus could restore GC compartmentalization and CSR, experimental *Cγ1-CDE, Foxo1*^*fl/stopRox*^, *R26-ZsGreen*^*stopRox*^ (in short *Foxo1*^*fl/stopRox*^) and control *Cγ1-CDE, Foxo1*^*fl/wt*^, *R26-ZsGreen*^*stopRox*^ (in short *Foxo1*^*fl/wt*^) mice were immunized with NP-CGG, subsequently treated with TAM (day 10-12 after immunization) and analyzed on day 14. The *R26-ZsGreen*^*stopRox*^ allele was included as an indirect reporter to identify Dre-recombined cells, i.e., FOXO1 re-expression in *Foxo1*^*stopRox*^ mice (Fig. 2C). Strikingly, TAM administration with subsequent FOXO1 re-activation in the *Foxo1*^*fl/stopRox*^ group completely restored the DZ compartment in ZsGreen^+^ GCBCs, up to the same levels as in *Foxo1*^*fl/wt*^ control mice (Fig. 2D). This observation also suggests a high efficiency of Dre-mediated recombination in this setting, since all rox sites available (here the rox sequences in *Foxo1*^*stopRox*^ and *R26-ZsGreen*^*stopRox*^ alleles) are targeted in the same cell. In contrast, ZsGreen^-^ GCBCs in the experimental *Foxo1*^*fl/stopRox*^ group, which had failed to recombine the *Foxo1*^*stopRox*^ allele by Dre and had remained a full FOXO1-KO, lost the physiological levels of the DZ/LZ ratio, at the expense of DZ cells (Fig. 2D) and in agreement with previous studies^14,15^. Additionally, CSR to IgG1 was robustly restored in ZsGreen^+^ GCBCs in the *Foxo1*^*fl/stopRox*^ group, albeit at a lower frequency compared to controls (Fig. 2E). We attribute this lower level to the fact that reporter ZsGreen^+^ cells from the *Foxo1*^*fl/stopRox*^ mice have resided in the GC only for a maximum of 4 days (after TAM administration), in contrast to the control cells, which have participated in the GC reaction from the beginning. This considerable rescue of class-switching in GCBCs at the peak of the GC reaction suggests that CSR can robustly be induced during the GC response, irrespective of its frequent occurrence at the pre-GCBC stage^13^.

**Figure 2.**
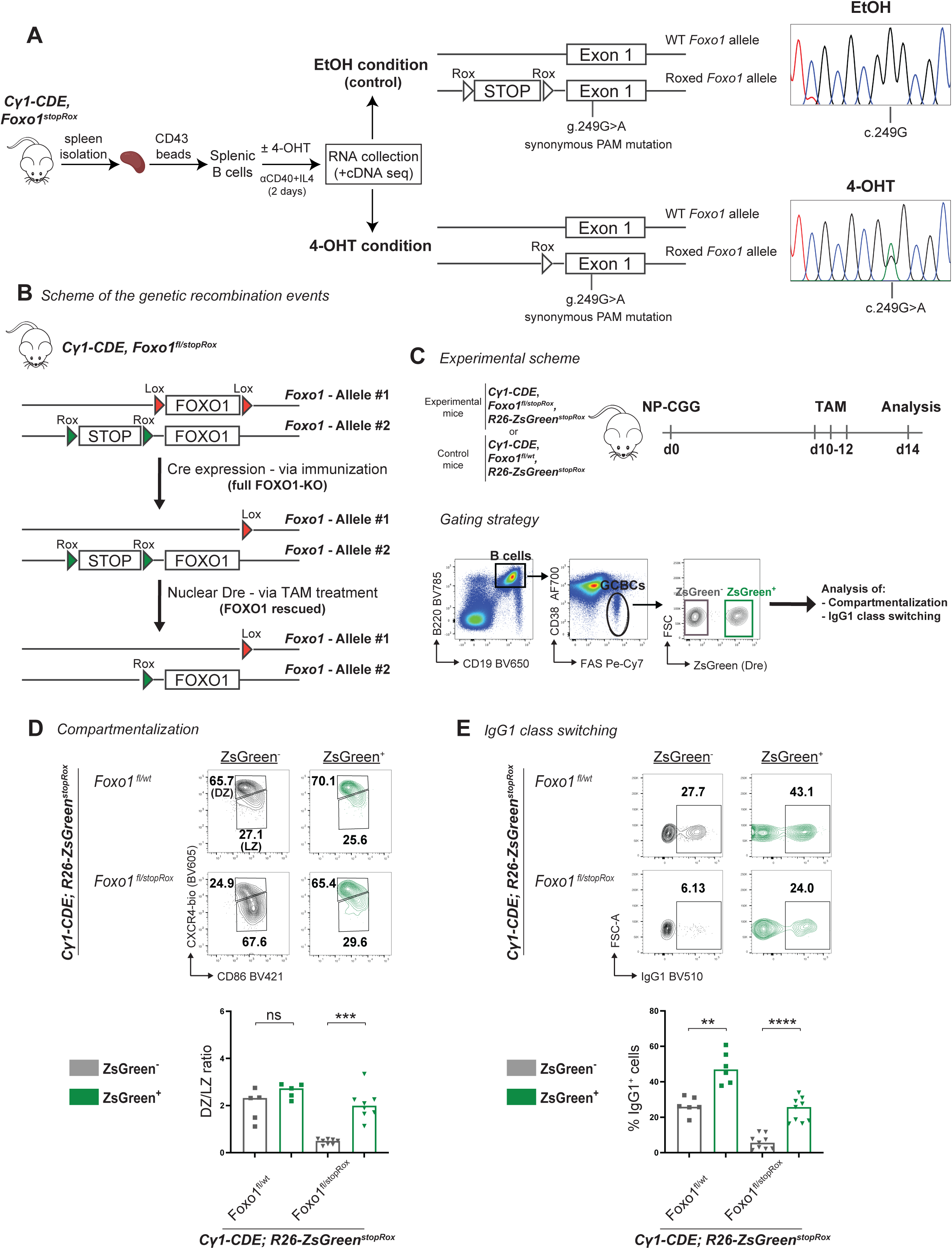
Sequential mutagenesis at the endogenous *Foxo1* locus. (A) Schematic representation of *in vitro* induction of the *Foxo1*^*stopRox*^ allele by *Cγ1-CDE* (left) and cDNA sequencing in activated *Cγ1-CDE, Foxo1*^*stopRox*^ splenic B cells (right). Expression of the targeted allele is observed only in the presence of 4-OHT, as demonstrated by the detection of the synonymous PAM mutation introduced during the targeting strategy of the transgenic strain. (B) Genetic scheme of recombination events after Cre and Dre activation at the *Foxo1* locus of *Foxo1*^*fl/stopRox*^ animals. (C) Experimental scheme (top) and representative gating strategy (bottom) employed for the phenotypic analysis of the GC compartment. (D-E) Top: Representative flow cytometry plots measuring the percentage of DZ and LZ (D) and IgG1^+^ (E) splenic GCBCs of mice of the indicated genotypes at day 14 after immunization. Bottom: Quantification of the DZ/LZ ratio (D) and the percentage of IgG1^+^ (E) ZsGreen^-^ (grey) and ZsGreen^+^ (green) splenic GCBCs. Statistics: Mann–Whitney test \*\**P* ≤ 0.01; \*\*\**P* ≤ 0.001; \*\*\*\**P* ≤ 0.0001; ns, not significant. Each symbol represents one mouse. Data from three independent experiments. Horizontal lines indicate the median.

In summary, our newly established *Cγ1-CDE* allele allows for highly efficient Cre and subsequent Dre recombination in the context of GCBCs. Thus, this tool can be used *in vivo* not only to model the sequential acquisition of genetic alterations acquired during malignant transformation but also to study the interplay of two or more genetic events or the function of a specific gene at different time points during the GC reaction. A similar strategy of sequential targeted mutagenesis could be applied in other cellular contexts.

## Abbreviations

4-OHT: 4-hydroxytamoxifen
BFP: blue fluorescent protein
Cγ1: immunoglobulin heavy constant gamma 1
CSR: class switch recombination
DZ: dark zone
GC: germinal center
GCBC: GC B cell;
KO: knockout
LZ: light zone
MBC: memory B cell
NP-CGG: 4-hydroxy-3-nitrophenylacetyl-conjugated chicken gamma globulin
PC: plasma cell
R26: Rosa 26
SHM: somatic hypermutation
TAM: tamoxifen.

## Acknowledgements

We thank all members of the K. Rajewsky lab for discussion, R. Kühn (MDC transgenic facility), H.P. Rahn (MDC FACS facility), G. Natale (K. Rajewsky lab, MDC) for excellent technical support and the MDC animal caretakers for their outstanding technical help. Figure schematics were created using images from BioRender. This work was supported by the European Research Council (ERC Advanced Grant 268921) to K.R., the Deutsche Krebshilfe (grant #70112800) to K.R. and M.J., and the Berlin School of Integrative Oncology (BSIO) to C.F.D. We apologize to colleagues whose work could not be cited owing to space limitation.

## Author contributions

C.F.D., E.K. and W.W. designed the experiments; C.F.D., E.K., W.W. and C.S. performed experiments; T.W. provided the *R26-ZsGreen*^*stopRox*^ strain; C.F.D. and E.K. analyzed data and wrote the manuscript; M.J. and K.R. supervised all aspects of the study, and reviewed and edited the manuscript.

