## Supplemental figure 1 + 2; material and methods for "*In vivo* sequential mutagenesis in germinal center B cells using a dual-recombinase approach: FOXO1 re-expression upon FOXO1 knockout rescues class switch recombination"

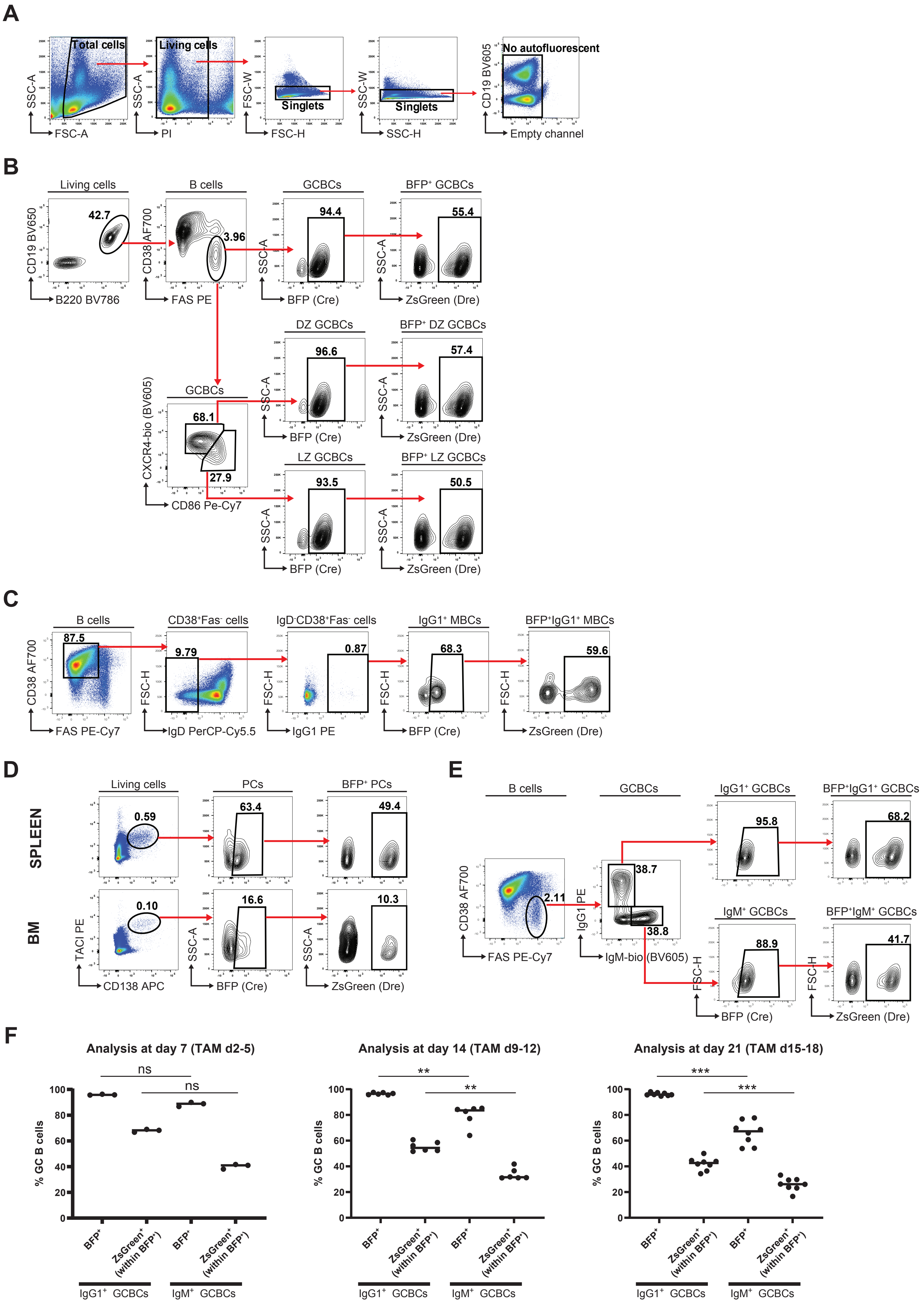

**Supplemental Figure 1. Characterization of the *Cγ1-CDE* strain.** (A-E) Representative gating strategies for non-autofluorescent living single cells (A) and reporter positive GCBCs (B) – including DZ and LZ GCBCs –; IgG1$^+$ MBCs (C); PCs (D) and IgG1$^+$/IgM$^+$ GCBCs (E). (F) Quantification of E at day 7 – TAM day 2-5 – (left panel), day 14 – TAM day 9-12 – (middle panel) and day 21 – TAM day 15-18 – (right panel) after NP-CGG immunization. Statistics: Mann-Whitney test $^{**}P \leq 0.01$; $^{***}P \leq 0.001$; ns, not significant. Each dot represents one mouse. Data from at least two independent experiments. Horizontal lines indicate the median.

### SUPPLEMENTAL FIGURE 2

**A**

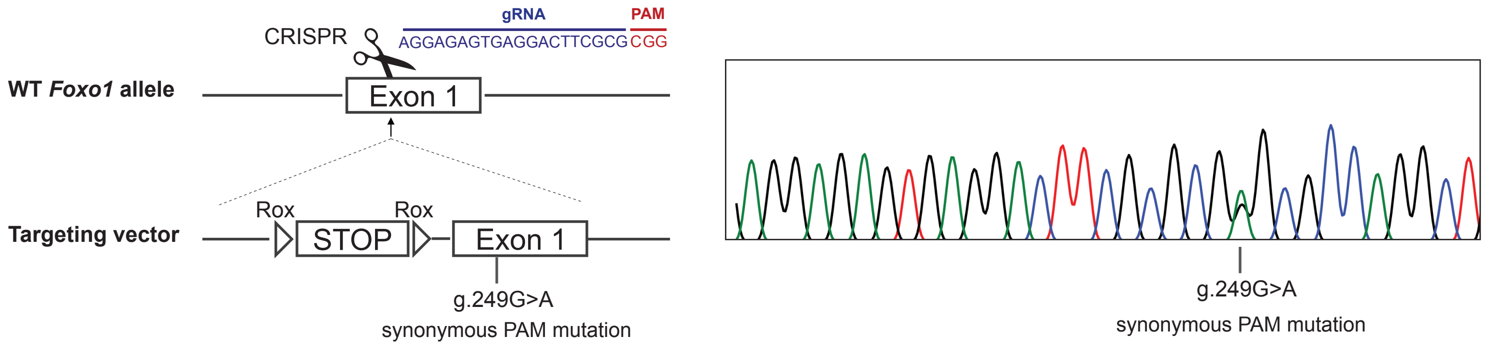

**Supplemental Figure 2. Targeting strategy at the endogenous *Foxo1* locus.** A STOP cassette flanked by rox sites was introduced in front of the endogenous Kozak sequence, and the sequence of exon 1 was modified in order to contain a synonymous PAM mutation in position g.449G>A which abrogated the genetic editing of the targeting vector used for the homologous recombination. The gRNA sequence used is shown in blue. Heterozygous mutant mice were verified by PCR and Sanger sequencing that allowed for the detection of the PAM mutation and thus verification of successful targeting.

**List of antibodies used for flow cytometry analyses**

| REAGENT | COMPANY | CLONE | CATALOG NUMBER |
|---|---|---|---|
| B220-BV786 | BioLegend | RA3-6B2 | 103246 |
| CD19-BV650 | BioLegend | 6D5 | 115541 |
| CD95/FAS-PE | BD Biosciences | JO2 | 554258 |
| CD38-AF700 | eBiosciences | 90 | 56-0381-82 |
| CD86-PE-Cy7 | BioLegend | GL-1 | 105014 |
| CXCR4-bio | Invitrogen | 2B11 | 1921475 |
| Streptavidin-BV605 | BioLegend | - | 405229 |
| IgG1-PE | BD Biosciences | A85-1 | 550083 |
| IgM-bio | eBiosciences | Il/41 | 13-5790-81 |
| CD86-BV421 | BioLegend | GL-1 | 105032 |
| IgG1-BV510 | BD Biosciences | A85-1 | 740121 |
| CD95/FAS-PE-Cy7 | BD Biosciences | JO2 | 557653 |

#### Mice, immunization, and TAM treatment

*Cγ1-cre, R26-BFP*$^{stopF}$ and *Foxo1*$^{fl}$ alleles have been described previously[1,2,3]. *R26-ZsGreen*$^{stopRox}$ is a derivative of the *R26-CAGS-lox-STOP-lox-rox-STOP-rox-ZsGreen* mice crossed to a *Deleter-Cre* line[4]. *Cγ1-CDE* and *Foxo1*$^{stopRox}$ strains were generated by CRISPR/Cas9-mediated homologous recombination in C57BL/6 mouse zygotes according to previously published protocols[5].

Mice were bred and maintained under specific pathogen-free conditions. 8- to 12-week-old male or female mice were immunized with 100 µg alum-precipitated NP-CGG (Ratio 10-19, LGC Biosearch Technologies Cat#N-5055B) intraperitoneally followed by TAM administration at the indicated time points (4 mg/day oral gavage, Tamoxifen, Sigma #T5648-5G, 99%). Experimental animal procedures were approved by the Landesamt für Gesundheit und Soziales Berlin (G0308/19, G0062/21).

#### Flow cytometry

Cells from spleen and bone marrow were collected in B cell medium (DMEM supplemented with 10% FCS, 1x NEAA, 1 mM sodium Pyruvate, 2 mM L-glutamine, 1 mM HEPES, 1x Penicillin-Streptomycin, 50 µM β-ME). Erythrocytes were lysed with Gey's solution and single cell suspensions were stained with antibody conjugates (see *List of antibodies used for flow cytometry analyses*) in PBS, pH 7.2, supplemented with 0.5% BSA and 2mM EDTA. Samples were analyzed on an LSRFortessa (BD BioSciences). Plots were generated using FlowJo software (BD FlowJo) and graphs by Prism software (GraphPad Prism).

#### B cell *in vitro* culture

Transgenic splenic B cells were isolated by CD43 depletion with magnetic anti-mouse CD43 microbeads (Miltenyi Biotech, Cat#130-049-081) according to the manufacturer's instructions. To remove the rox-flanked STOP cassette, B cells were activated with IL-4 (25 ng/mL, R&D, Cat#404-ML) and αCD40 (1-2 µg/mL, BioLegend, HM40-3, Cat#102908) and cultured in the presence of 4-OHT (1 µM, Sigma Aldrich, Cat#H6278). Three days later total RNA was extracted using the AllPrep DNA/RNA Mini Kit (Qiagen, #80204) according to the manufacturer's instructions. For cDNA synthesis 500 ng of RNA were used per reaction and reverse transcription was performed with SuperScript$^{TM}$ II Reverse Transcriptase (Invitrogen, #18064014). A mixture of random and oligo(dT) primers was used following the manufacturer's protocol. PCR amplification of *Foxo1* cDNA was performed using the following primers: forward 5'-ATGGCCGAAGCGCCCCAGGTGGTGGAGAC-3' and reverse 5'-CCTACTTCAAGGATAAGGGCGACAGCAAC-3'. PCR products were gel purified using the NucleoSpin Gel and PCR clean-up kit (Macherey and Nagel, #740609.250) and sequenced by Sanger Sequencing.

#### Supplementary information references:
